# The influence of tension-compression switches on brain anisotropic modelling

**DOI:** 10.64898/2026.04.10.717701

**Authors:** Chengbin Li, Zhou Zhou

## Abstract

Finite element (FE) head models are valuable tools for investigating brain injury mechanics, with their reliability critically dependent on accurate material modelling. White matter (WM) is often considered mechanically anisotropic due to its aligned axonal fiber architecture and is commonly represented using fiber-reinforced hyperelastic formulations such as the Gasser-Ogden-Holzapfel (GOH) model. A fundamental assumption of the GOH model is that fibers contribute only in tension and not in compression, requiring the use of tension-compression switches. However, inconsistencies were noted in the formulation of tension-compression switches with the influence on computational biomechanics unknown. To address this knowledge gap, three commonly used switching schemes - differing in both the switching parameter and the treatment of compressed fibers - were theoretically elaborated and numerical implementation within the GOH framework to simulate the mechanical anisotropy of WM in impact simulations. Results from the case-based and group-level analyses demonstrated that both the switching parameter and the treatment of compressed fibers affected WM deformation. Significant cross-scheme strain differences were noted in the first principal strain at the element level and fiber strain at the fiber level. These findings highlighted the mechanical role of tension-compression switch in the GOH-based brain modelling and advocated the adoption of fiber stretch itself as the switching parameter to discriminate the tensile and compressive fibers. The current study provides important guidance for the anisotropic constitutive models in brain tissue and calls for direct verification of the tension-compression switch hypothesis in axonal fibers.

## 1. Introduction

Traumatic brain injury (TBI) is a leading cause of death and disability and poses a substantial public health burden. Globally, epidemiologic data showed that about half of the world’s population experienced at least one TBI over their lifetime, and the annual economic cost related to TBI was around 400 billion dollars [1]. In the United States, approximately 60600 TBI-related deaths occurred in 2019 [2]. In Europe, more than 2 million people are admitted to hospital each year because of TBI, with an incidence rate of 287/100000 and about 82000 deaths [3]. To address this urgency, efforts to advance fundamental understandings of brain injury biomechanics are crucial for developing more effective prevention strategies.

Finite element (FE) head models are valuable tools to quantitatively investigate the mechanics of TBI. As numerical surrogates, FE head models can offer spatiotemporal detailed information of intracranial responses secondary to external impacts of varying magnitude and direction. Such valuable information is otherwise challenging or impossible to attain due to enormous expenses and technical difficulties (e.g., highly controlled cadaver experiments) or ethical reasons (e.g., imposing injury-level accelerations on living humans) [4, 5]. Modern FE head models are featured with detailed representations of brain anatomy [6, 7], advanced constitutive descriptions to capture nonlinear features of brain material with tension-compression asymmetry and rate dependency [8, 9], and fluid-structure interaction representations of the brain-skull interface [10, 11] and brain-ventricle interface [6, 12]. In recent years, the FE-predicted responses started to show promising correlations with experimentally quantified pathology in animals (e.g., pig [13] and rat [14]) and clinically diagnosed injury in humans (e.g., concussion [15], subdural hematoma [16]).

Accurate brain tissue modeling is essential for ensuring the biofidelity of FE head models. A major challenge in this task arises from the complexity of the brain, which is composed of multiple substructures marked by different mechanical properties. The white matter (WM) has long been known to be an anisotropic structure, consisting of bundles of axonal fibers with preferential orientations that connect different regions of the brain. With the advance of imaging techniques, definitive information on WM fiber orientation and distribution of large bundles can now be derived from diffusion tensor imaging (DTI) [17]. In parallel, several material testing studies have reported anisotropic behavior in brain tissue, especially for some WM regions that the stiffness parallel to the fiber tracts was larger than that perpendicular to them [18-25]. These converging findings motivate the use of anisotropic constitutive models that incorporate fiber orientation to represent WM material. Indeed, incorporating fiber-reinforced anisotropy informed by DTI for the brain tissue modelling has been reported by several independent groups to improve the model validation performance [26-29] and the accuracy of injury prediction [13, 28, 30-33].

Among available anisotropic formulations, the constitutive model proposed by Gasser, Ogden, and Holzapfel (GOH) [34] has been widely adopted in brain anisotropic modelling [13, 26-28, 30, 32, 35-42], compared with other alternatives, such as the Puso-Weiss model [33, 43, 44] and the quadratic reinforcing model [45]. This might be related to the fiber-reinforcement dispersion parameter (*κ*, as detailed in section 2.2) in the GOH model, capable of accounting for the degree of anisotropy owing to different axonal alignment [40]. This GOH model has also been successful in capturing the mechanical responses of other biological soft tissues (e.g., artery [34], skin [46], tendon [47]) and is widely recognized as one of the most important contributions to the biomechanics [48].

One central hypothesis in the GOH model is that collagen fiber (axonal fibers in the context of brain tissue) could not support any compression and would buckle under the smallest compressive load [34]. This means that, in tension, the axonal fiber leads to a stiffening response. Under compression, however, the axonal fiber stores very little or no energy, and the matrix bears the compressive load [49]. The numerical implementation of this assumption requires a tension-compression switch to exclude compressed fibers, which is also essential for reasons of stability [34, 50]. Nevertheless, there is currently no consensus on how this switch is formulated. For example, studies adopted either the fourth invariant (*Ī*_4*α*_) or a strain-like measure (*Ē*_*α*_, with the exact definition in section 2.2) as the switching parameter [27, 32, 35, 39], despite that the latter one has been widely challenged in terms of its capability of distinguishing between tensile and compressive fiber states [27]. In addition, most implementations for WM modelling fully eliminate the contribution of compressed fibers, deviating from the original GOH formulation [34], which retains an isotropic contribution. These inconsistencies raise two important questions: 1) How sensitive is the brain response to the choice of switching parameter (i.e., *Ī*_4*α*_ vs. *Ē*_*α*_) 2) How do different treatments of compressed fibers influence model behavior?

The present study aimed to investigate the mechanical influence of tension-compression switch formulations within the GOH framework for anisotropic brain tissue modelling. Three candidate switching schemes, differing in both the switching parameter and the treatment of compressed fibers, were implemented in an FE head model to represent WM as a fiber-reinforced anisotropic material. Their effects were evaluated by comparing brain responses during head impacts. It was hypothesized that the tension-compression switch schemes affected WM deformation. This study clarified the ambiguity of tension-compression switches in the GOH model and provided important guidance for their implementation of brain tissue modelling.

## 2. Method

### 2.1 Finite element head model

The FE head model used in the current study was previously developed at the Kungliga Tekniska högskolan by Kleiven [8] using the LS-DYNA software. The model included the scalp, skull, brain, subarachnoid cerebrospinal fluid, ventricles, dura mater, falx, tentorium, pia mater, 11 pairs of the largest parasagittal bridging veins, and a simplified neck with the extension of the spinal cord (Fig 1). The material properties of the FE model are summarized in Table 1, except for the brain that is elaborated in Sections 2.2 and 2.3. A penalty-based contact was defined between the brain and skull that permitted sliding in the tangential direction and delivered tension and compression in the radial direction. The model has been validated against experiments of brain-skull relative motion [11, 51], intracranial pressure [51], and brain strain [52]. It has been used to study the brain injury risk in real-life trauma accidents (e.g., bicycle accidents [53], lacrosse [54], high-school American footballs [55]), assess the protective efficacy of safety products (e.g., bicycle helmet [56], impact-absorbing pavement [57], face airbag [58]), and evaluate and improve the method for headgear homologation (e.g., designing the oblique impact helmet testing method via virtual simulations [59-61]).

**Table 1.** Material properties of the FE head model. Note that *K* represents the bulk modulus, EA represents the force per unit strain, and GOH material represents the constitutive law proposed by Gasser, Ogden, and Holzapfel as detailed in section 2.2.

| Tissue | Young's modulus (MPa) | Density (kg/dm<sup>3</sup>) | Poisson's ratio |
| --- | --- | --- | --- |
| Outer compact bone | 15000 | 2.00 | 0.22 |
| Inner compact bone | 15000 | 2.00 | 0.22 |
| Porous bone | 1000 | 1.30 | 0.24 |
| Neck bone | 1000 | 1.30 | 0.24 |
| Brain | Hyperelastic fiber-reinforced GOH material (Table 2) |  |  |
| Cerebrospinal fluid | $K = 2.1$ GPa | 1.00 | |
| Sinuses | $K = 2.1$ GPa | 1.00 | |
| Dura mater | 31.5 | 1.13 | 0.45 |
| Falx/Tentorium | 31.5 | 1.13 | 0.45 |
| Pia mater | 11.5 | 1.13 | 0.45 |
| Bridging veins | $EA = 1.9$ N | | |

**Fig 1.**
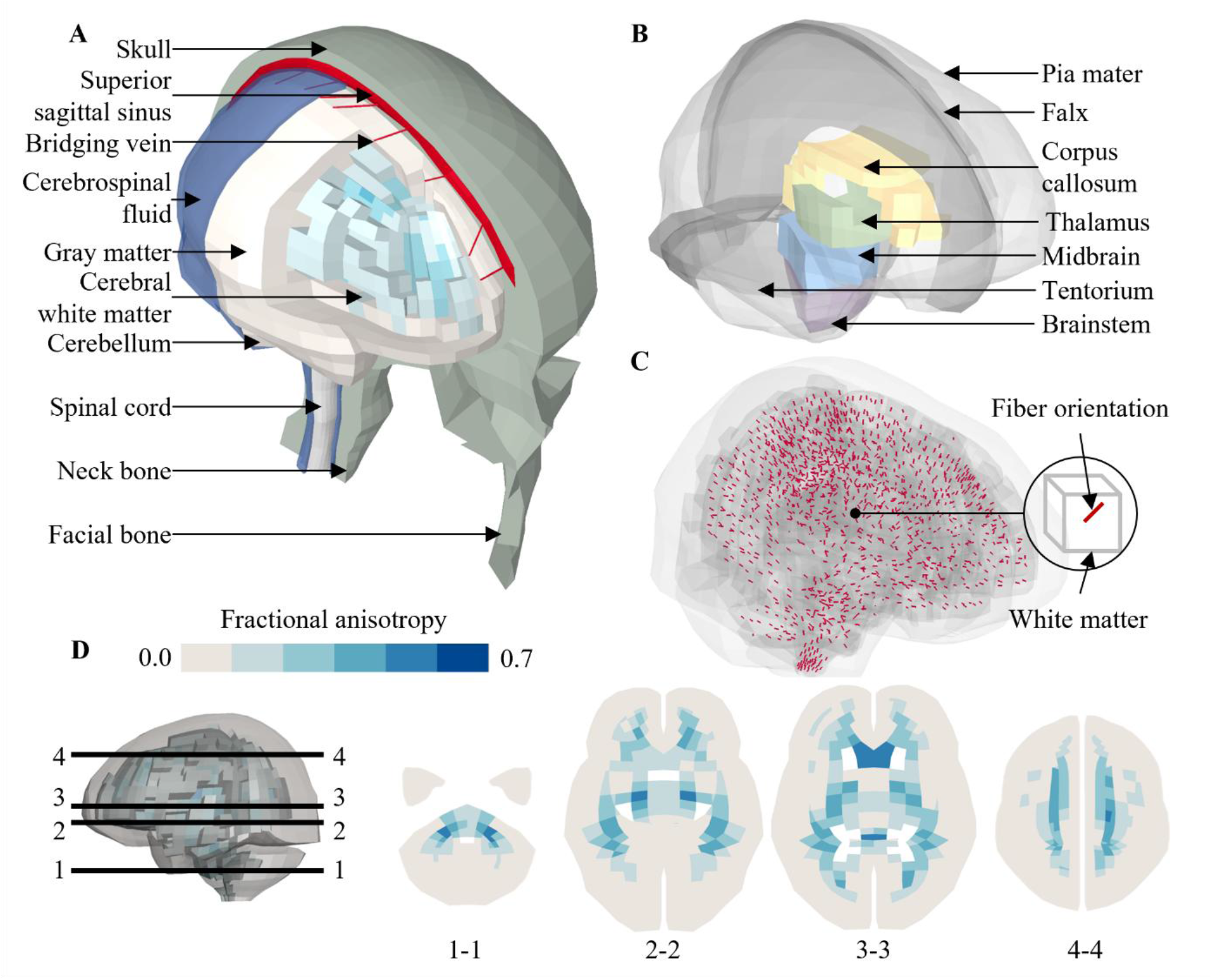
(A) Isometric view of the finite element model of the human head; (B) Isometric view of membranes and deep brain structures; (C) Isometric view of the finite element model with embeded fiber orientation (red) for anisotropic material modelling of brain tissue; (D) Axial cross-sections of fractional anisotropy contour of the white matter elements in the FE model.

### 2.2 Brain anisotropic material modelling

To inform the anisotropic brain material modelling, the degree of anisotropy and fiber orientation delineated by DTI were coupled with the FE model on an element basis. Specifically, the brain mesh of the head model was voxelized to generate a reference volume, which was aligned to the volume of the ICBM DTI-81 atlas [17] via an affine registration. For each brain element in the FE model, corresponding voxels in the DTI atlas exhibiting spatial alignment with the given WM element were identified. The diffusion tensors of these identified voxels were averaged to extract the mean anisotropic information, including the degree of anisotropy [characterized by the fractional anisotropy (FA) value] and fiber orientation (i.e., characterizes by one vector with unit length (***n***_0*α*_), Fig 1C). Those elements with an FA value over 0.2 were considered as WM elements (N = 1197, Fig 1D), and the ***n***_0*α*_ of each WM element was defined via the *ELEMENT_SOLID_ORTHO keyword in LS-DYNA input deck. Further details regarding the coupling between DTI and FE brain model are available in previous studies by Giordano, et al. [39] and Zhou, et al. [62].

The brain tissue was modelled using the hyper-elastic fiber-reinforced anisotropic material using the formulation proposed by Gasser, Ogden, and Holzapfel (GOH) [34] to account for the WM anisotropy. In this material formulation, the strain energy function ***Ψ*** was defined as follows, based on the incompressible assumption of the brain tissue:

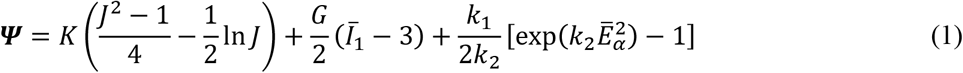

where *K* represents the bulk modulus, *J* is equal to the determinate of the deformation gradient and closer to 1, *G* represents the shear modulus, 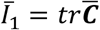 is the first invariant of the isochoric right Cauchy-Green strain tensor 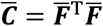 in which 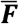 is the isochoric deformation gradient, *k*_1_ and *k*_2_ describe the fiber stiffness. *Ē*_*α*_ = *k*(*Ī*_1_ − 3) + (1 − 3*κ*)(*Ī*_4*α*_ − 1) is a Green-Lagrange strain-like quantity. *Ī*_4*α*_ is a tensor invariant equal to the square of the isochoric stretch in the direction of the mean orientation ***n***_*0α*_ of the family of fibers, where 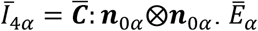 was used to calculate the fiber contribution, which was simplified to be linear as *k*_2_ →0 [36, 37].

In the current study, the dimensionless structure parameter *κ* accounting for the orientation dispersion of the axonal fiber was related by the FA values by assuming similarity between the mechanical and diffusion anisotropy (Table 2B) [40, 42]. The gray matter was assumed to be isotropic (*κ* = 0.3333), and the WM elements were binned into five groups based on the FA values (Fig 1C and Table 2B), similar to the approach of Giordano and Kleiven [40]. The exact value of brain material properties is summarized in Table 2, the same as those in Giordano and Kleiven [28]. Note that the viscoelastic behavior was not included in this study, ensuring a focused investigation of the effect of tension-compression switches.

**Table 2.** (A) Properties used in the brain tissue; (B) Discretization of fractional anisotropy in intervals and respective *κ* value for FE simulations. Note that G is shear modulus, K is bulk modulus, *k*_1_, *k*_2_ describe the fiber stiffness and *κ* is dimensionless structure parameter.

| (A) | Parameter | Value | (B) | FA range | $\kappa$ value |
| --- | --- | --- | --- | --- | --- |
|  | G (Pa) | 2990 |  | 0.0~0.2 | 0.3333 |
|  | K (MPa) | 50 |  | 0.2~0.3 | 0.2732 |
| | $k_1$ (Pa) | | | 0.3~0.4 | 0.2500 |
|  | Occipital lobe | 31672 |  | 0.4~0.5 | 0.2273 |
|  | Frontal and parietal lobe | 30242 |  | 0.5~0.6 | 0.2000 |
|  | Thalamus | 37097 |  | 0.6~0.7 | 0.1667 |
|  | Corona radiata | 37143 |  |  |  |
|  | Corpus callosum | 33709 |  |  |  |
| | $k_2$ | $\rightarrow 0$ | | | |

### 2.3 Implementation of tension-compression switches in GOH model

To analyze the influence of tension-compression switch in the GOH model, three different candidate schemes were implemented. These schemes were either used in the literature or available via commonly used FE softwares. Note that, when the fiber was deemed in tension, the three schemes (equations 2-4) were identical with*Ē*_*α*_ = *κ*(*Ī*_1_ − 3) + (1 − 3*κ*)(*Ī*_4*α*_ − 1). Fundamental differences existed among the three schemes in terms of the switching parameters and the treatment of fiber in compression.

Scheme 1 (Equation 2) employed *Ē*_*α*_ as the switching parameter. When *Ē*_α_ *≤* 0 (i.e., the fiber was in compression), *Ē*_*α*_ became zero, denoting that the compressed fiber had no mechanical contribution. Scheme 1 was implemented in the Abaqus software [63] and FEBio software [64] and was the most commonly used scheme for the anisotropic modelling of brain tissue [26, 28, 35-37, 39-42].

Scheme 2 (Equation 3) instead used *Ī*_4*α*_ as the switching parameter. When *Ī*_4*α*_ ≤ 1 (i.e., the fiber was in compression), *Ē*_*α*_ was assigned to be zero, the same as that in scheme 1. Scheme 2 was similarly implemented in LS-DYNA software [65, 66] and ANSYS software [67] and was used to model the anisotropic mechanical behavior of axonal fiber tracts in the WM [13, 27] via user-defined subroutines.

Scheme 3 (Equation 4) also adopted *Ī*_4*α*_ as the switching parameter. When the fiber was in compression (i.e., *Ī*_4*α*_*≤*1), *Ē*_*α*_ reduced to *κ*(*Ī*_1_ − 3), thereby retaining an isotropic contribution. Scheme 3 was originally proposed by Gasser, et al. [34], although it has been interpreted differently in the literature, leading to the alternative implementations represented by schemes 1 and 2.

**Scheme 1:**

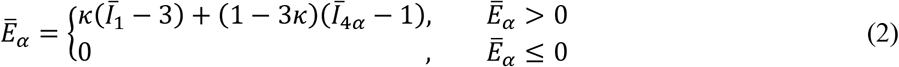

**Scheme 2:**

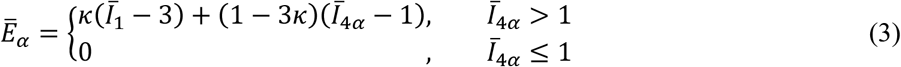

**Scheme 3:**

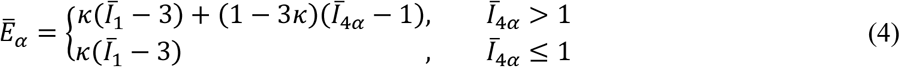

To illustrate the differences among the three tension-compression schemes, the *Ī*_4*α*_-*Ē*_*α*_ relationship are plotted in Fig 2 in the case of uniaxial tension and compression with the loading direction along the fiber orientation. In the current study, all three schemes were implemented in LS-DYNA via use-defined material.

**Fig 2.**
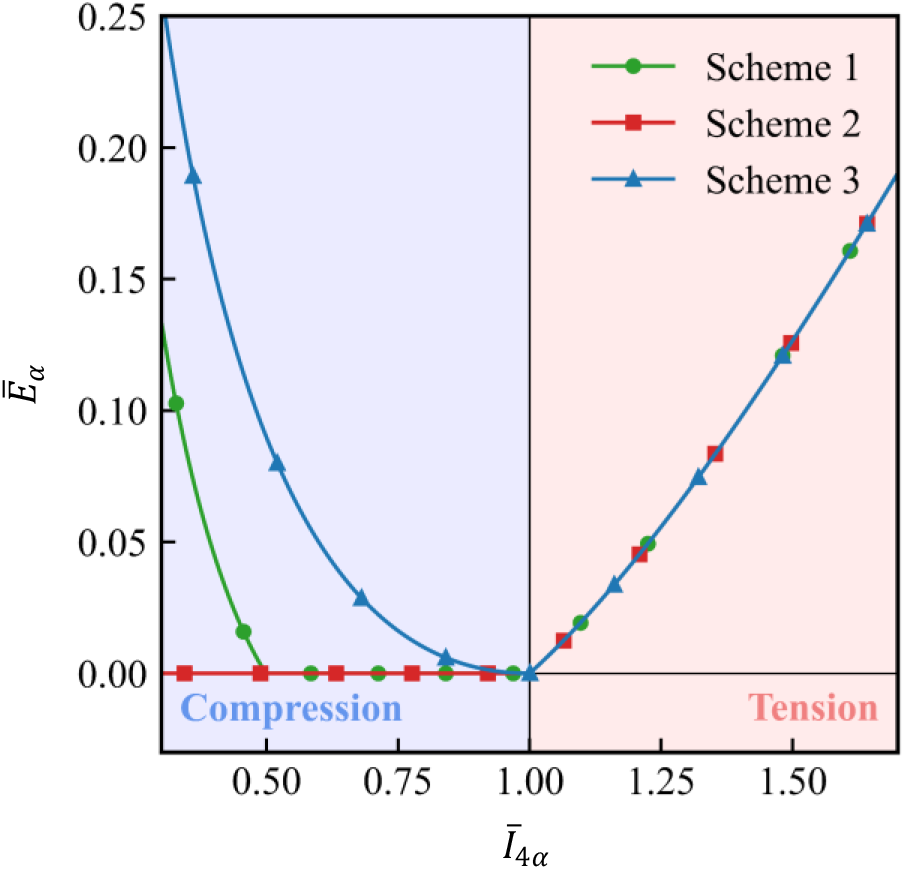
The plot of the function *Ē*_4*α*_ in relationship to *Ī*_4*α*_ in the three candidate schemes under uniaxial tension and compression with the loading direction along the fiber orientation. For illustrative purpose, we defined 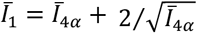 with the incompressibility constraint (i.e., *J =*1) and *κ* = 0.2732. It can be noted that *Ē*_*α*_ > 0 does not necessarily indicate tensile fiber (i.e., *Ī*_4*α*_ > 1), indicating that the *Ē*_*α*_ is an inappripriate proxy of fiber stretch.

### 2.3 Loading condition

To analyze the influence of tension-compression switches on brain responses, whole-head impact simulations were performed. Two concussive impacts measured by instrument mouthguards from collegiate football players were included: one resulting in loss of consciousness (Fig 3D) and the other self-reported with post-concussive symptoms [68]. In addition, 36 impacts (15 concussed and 21 non-concussed) from the National Football League were analyzed. The kinematic data for these professional football impacts were initially obtained from laboratory reconstruction using Hybrid III anthropometric test dummies [69-71] and recently corrected by Sanchez, et al. [72]. Among the 53 reconstructed cases, only 36 satisfied the requirement that the maximum strains occur within the available impact duration and were therefore retained for subsequent analysis, the same as the strategy used by Zhou, et al. [73]. These 38 impacts spanned a range of impact planes and loading severities (Fig 3) and were associated with different injury outcomes.

**Fig 3.**
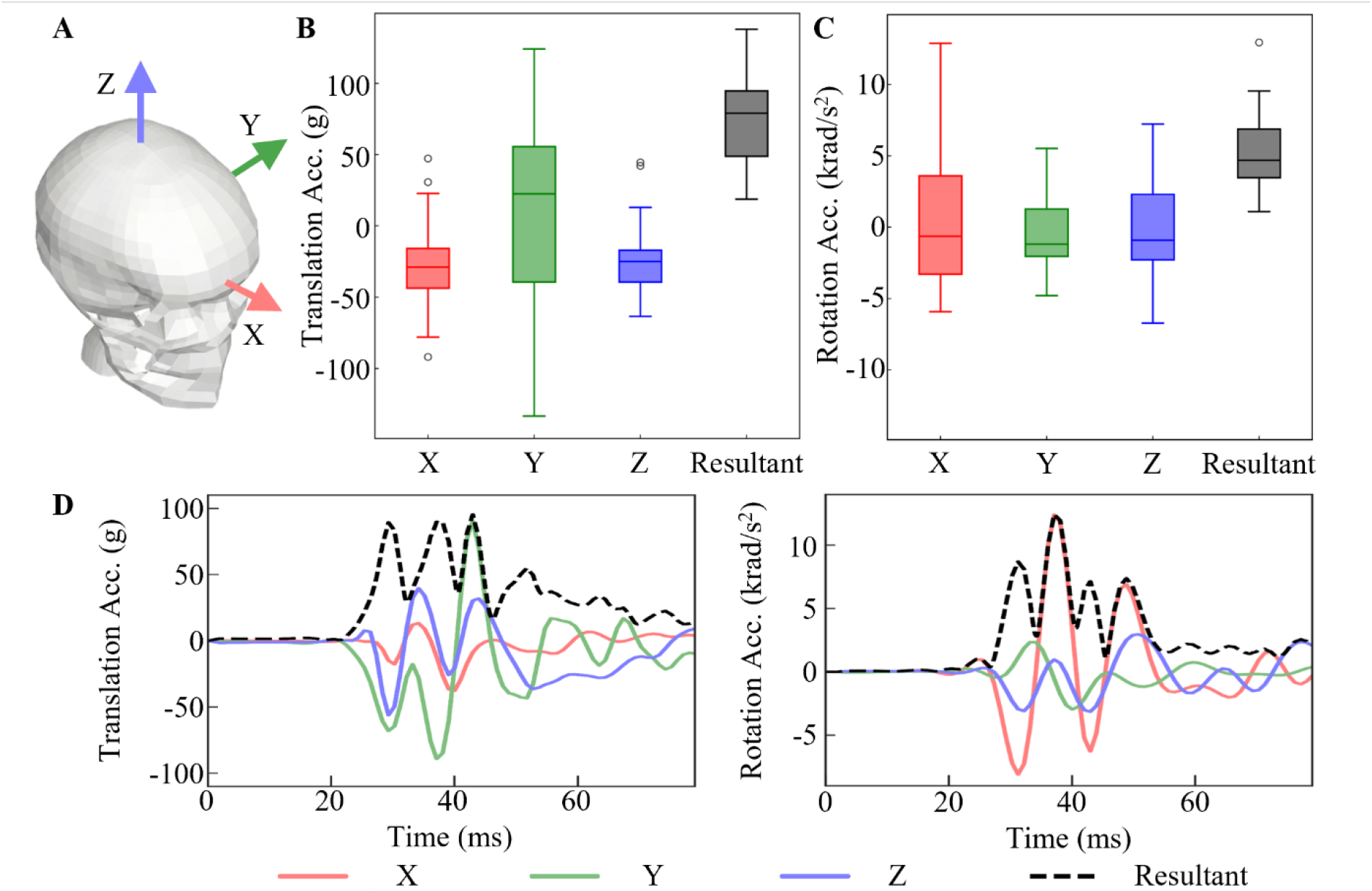
(A) Isometric view of the finite element model and a skull-fixed coordinate system and corresponding axes are illustrated with the origin at head’s center of gravity; (B) Boxplot of translation acceleration peaks of the 38 simulated cases; (C) Boxplot of rotation acceleration peaks of the 38 simulated cases; (D) Head model loading condition for the illustrative case. Note that, for each box plot in subfigures B and C, the central line is the median value, and the upper and lower edges of the box are the 25th and 75^th^ percentile values, while the outliers are shown as ‘‘o’’ symbol outside the box.

For each simulation, the six directional impact pulses were prescribed to a node located at the center of gravity of the FE head model and this node was attached to the rigid skull. All simulations were solved by LS-DYNA (versions 16.0 double precision). For a simulation of 80 ms, it took 40 minutes to solve using 16 central processing units on Linux.

### 2.4 Data analysis

Two strain-based metrics were calculated from WM elements (as was the region of interest in the current study) to quantify the potential influence of tension-compression switch schemes. The first principal strain, characterizing the maximum stretch within the WM element, was calculated as the maximum eigenvalue of the Green-Lagrange strain tensor. The fiber strain was computed as 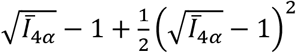 to quantify the isochoric normal Green-Lagrange strain along the fiber direction. The time-accumulated peaks of the first principal strain and fiber strain were termed as maximum principal strain (MPS) and maximum fiber strain (MFS), respectively. For each head impact, the MPS and MFS across all WM elements were identified, representing the most severe deformation at the element and fiber level, respectively.

While the three tension-compression switch schemes were theoretically different (Equations 2-4 and Fig 2), the interdependence between the strain response and tension-compression switch schemes within the context of brain computational biomechanics remained unclear. To assess scheme-dependent differences in the overall distributions of MPS and MFS across impacts (N = 38), the Friedman test was applied to evaluate overall differences among schemes, followed by the Nemenyi post-hoc test to identify specific pairs with the difference being significant. All the statistical analyses were performed using custom scripts in Python (v 3.11.8).

## 3. Results

### 3.1 Case illustration on the influence of tension-compression switches

To demonstrate the sensitivity of brain responses on the tension-compression switch scheme, one concussive impact (i.e., the one in Fig 3D) was employed as an illustrative case. The time history curves of the first principal strain of one randomly selected element (*κ* = 0.2500) are shown in Fig 4A. For the two schemes that differed in switch parameters (i.e., *Ī*_4*α*_ in scheme 1 vs. *Ē*_*α*_ in scheme 2), the strain magnitude in scheme 2 was much higher than that in scheme 1, resulting in an inter-scheme MPS difference of 25.7%. For the two schemes differed in the treatment of compressed fibers (i.e., 0 in scheme 2 vs. *κ*(*Ī*_1_ − 3) in scheme 3), scheme 2 yielded a MPS value that was 33.8% higher than that in scheme 3. When examining the response of fiber strain (Fig 4B) of another element (*κ* = 0.2732), scheme 2 predicted the highest fiber strain with the MFS value as 0.405, while schemes 1 and 3 estimated the MFS values as 0.302 and 0.282, respectively.

**Fig 4.**
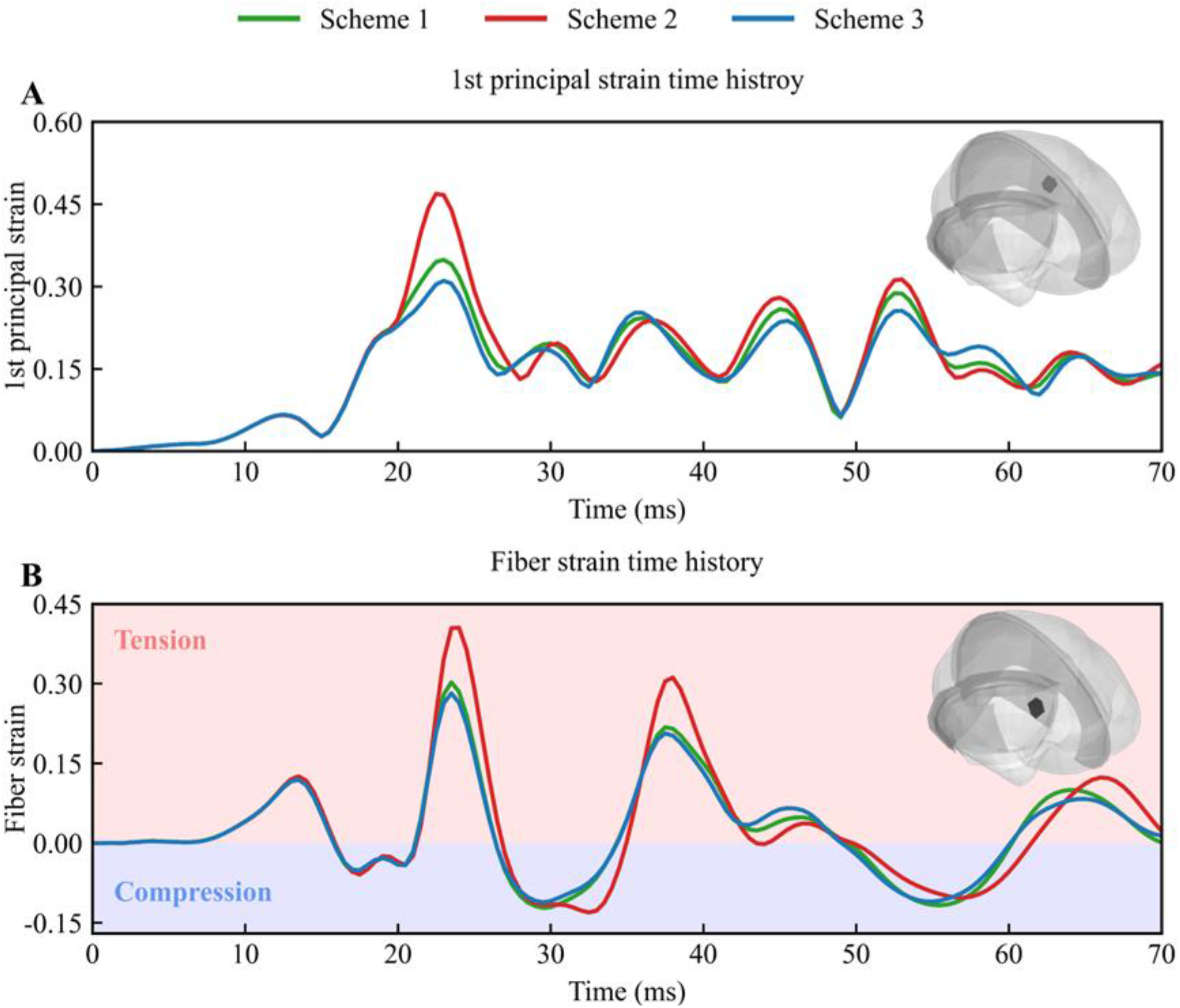
Influence of three tension-compression switch schemes on strain time history responses in one concussive impact. (A) Comparisons of time history curves of the first principal strain for one element in the cerebral white matter; (B) Comparisons of time history curves of the fiber strain for one element in the thalamus.

To quantify how the fiber loading modes varied with the tension-compression switches, WM elements with fibers in compression (Fig 5) were identified at a representative time point (i.e., 27 ms) for the illustrative impact. When using *Ē*_*α*_ as the switch parameter (i.e., scheme 1), fibers in 218 WM elements were in compression. When using *Ī*_4*α*_ as the switch parameter, the number of WM elements with compressed fibers were 189 for scheme 2, and 183 for scheme 3. The spatial distribution of fiber-compressed WM elements in scheme 1 differed notably from those in schemes 2 and 3. For example, scheme 1 predicted much fewer WM elements with the fibers in compression in the brainstem compared to those in schemes 2 and 3 (as highlighted by the ellipses in Fig 5). In contrast, the fiber-compressed WM element identified by schemes 2 and 3 were largely overlapped with minor differences noted. For example, an arrow in the upper-middle subfigure of Fig. 5 pointed to a WM element with the fiber in compression that was uniquely identified by scheme 2.

**Fig 5.**
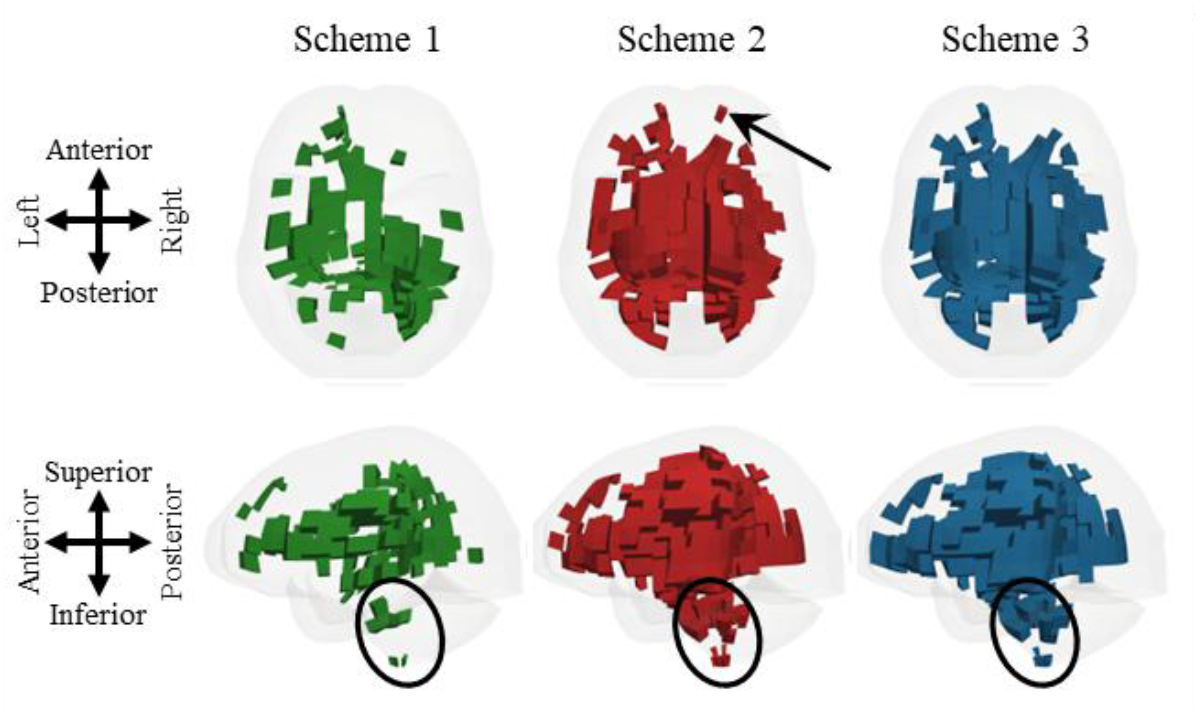
Top and side view of the WM elements with the embedded fiber in compression based on the three tension-compression switches at time of 27 ms of the illustraive case. Note that the arrow in the upper-middle subfigure pointed one WM element in which the embedded fiber were in compression only by scheme 2, while the ellipses in the low row highlighted the cross-schemes difference of fiber-compressed WM elements in the brainstem.

To evaluate the influence of tension-compression switches on deformation responses of WM elements, the results in scheme 2 were used as the baseline to calculate the absolute value of relative strain difference with respect to schemes 1 and 3 (Fig 6). For the comparison between schemes 1 and 2 (different switching parameters but identical treatment of compressed fibers), the maximum relative differences in absolute value were up to 0.334 for MPS, and up to 0.103 for MFS. When comparing schemes 3 and 2 (the same switching parameter but differ in the treatment of compressed fibers), the maximum absolute values of relative differences were 0.474 for MPS, and 0.124 for MFS.

**Fig 6.**
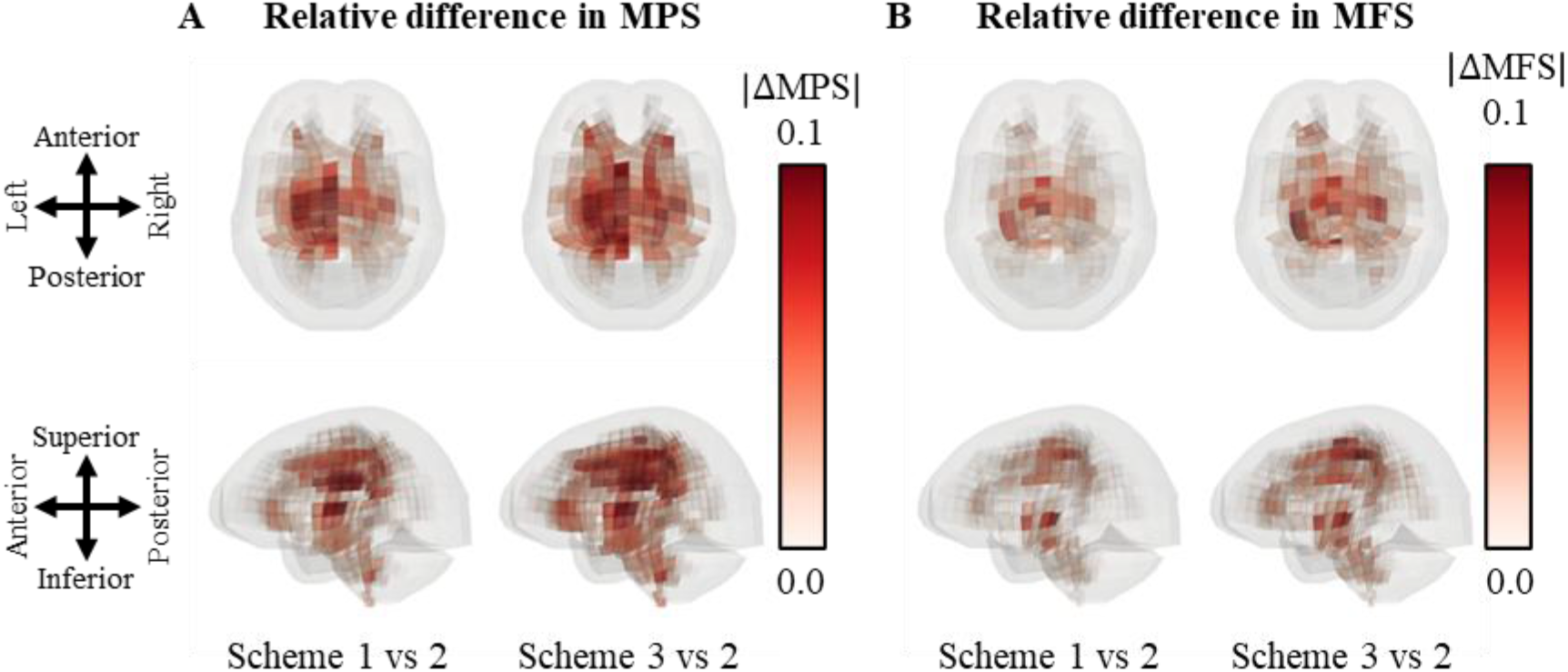
Top and side view of the brain showing the influence of three tension-compression switch schemes on the element-wise strain difference (in the form of absolute value) based on MPS and MFS in one illustrative case. (A) Absolute value of relative differences in MPS between schemes 1 and 2 (left) and between scheme 2 and 3 (right); (B) Absolute value of relative differences in MFS between schemes 1 and 2 (left) and between scheme 2 and 3 (right).

### 3.2 Group analysis on the influence of tension-compression switches

To evaluate the influence of tension-compression switch at the group level, MPS and MFS peaks across the entire WM were analyzed across all 38 simulated impacts. For the MPS (Fig 7A), significant differences were observed between schemes 1 and 2, as well as between schemes 2 and 3. For the MFS (Fig 7B), only schemes 1 and 2 showed statistically significant differences.

**Fig 7.**
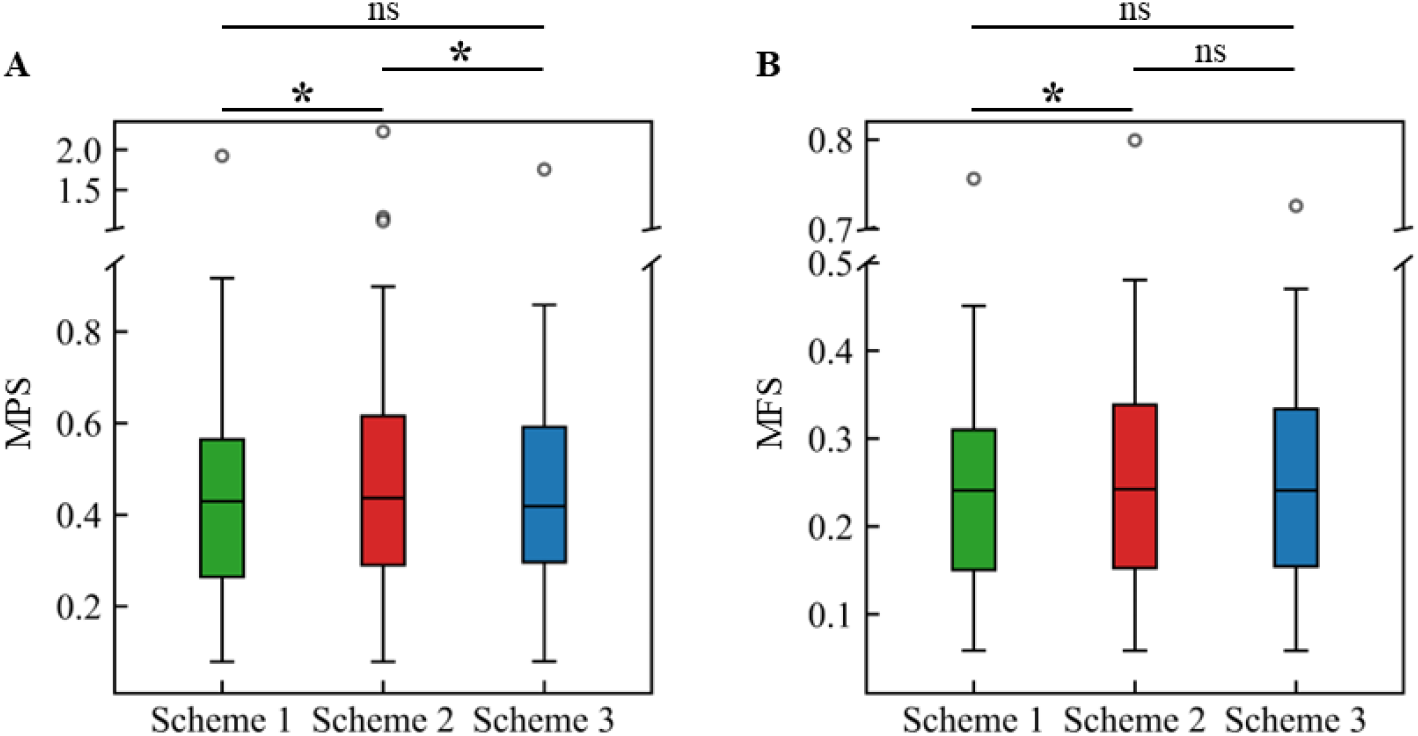
Influence of tension-compression switch schemes on MPS and MFS. The values (N=38) are shown in green for scheme1, red for scheme 2, and blue for scheme 3. (A) Comparisons of MPS; (B) Comparisons of MFS. On each box, the central line is the median value, and the upper and lower edges of the box are the 25th and 75^th^ percentile values, while the outliers are shown as ‘‘o’’ symbol outside the box. Statistical differences in Nemenyi post-hoc test are reported (ns: not significant; * indicates the paired comparison meets the significant level for difference).

## 4. Discussion

The current study presented a theoretical elaboration and numerical implementation of three candidate tension-compression switches (Equations 2-4) in the GOH model to simulate the mechanical anisotropy of WM in impact simulations. The results showed that both the switching parameter (*Ē*_*α*_ vs. *Ī*_4*α*_) and the treatment of fiber contribution in compression (0 vs. *κ*(*Ī*_1_ − 3)) influenced the MPS at the element level and MFS at the fiber level, verifying the hypothesis that the tension-compression switch affected the WM deformation. This study has important implications for anisotropic brain modelling by: 1) elucidating the theoretical differences and biomechanical consequences of commonly used tension-compression schemes for axonal fiber modelling; 2) pinpointing that the Green-Lagrange strain-like quantity *Ē*_*α*_, as is implemented in the Abaqus and FEBio softwares and widely used by the brain modeler, is not an appropriate proxy of fiber stretch (*Ī*_4*α*_).

The current study underscored the mechanical role of tension-compression switch in the GOH model, generally aligning with several earlier works focused on continuum theory. For example, Holzapfel and Ogden [74] and several their subsequent studies [49, 75] mathematically clarified two switching criteria - one proposed by Gasser, et al. [34] and the other implemented in Abaqus software (i.e., scheme 3 and scheme 1 in the current study) - and concluded that *Ē*_*α*_ used in scheme 1 was an inappropriate switching parameter. These two schemes were also evaluated by Latorre and Montáns (2016), who concluded that the stress responses were significantly different for different switching criteria. The current work extended the previous comparison by incorporating an additional switching scheme (i.e., scheme 2) which was more recently implemented in two widely used FE softwares, i.e., LS-DYNA and ANSYS. This inclusion enabled the new finding that, in addition to the switching parameters (*Ē*_*α*_ vs. *Ī*_4*α*_), the treatment of fiber’s contribution in compression (i.e., 0 in scheme 2 vs. *κ*(*Ī*_1_ − 3) in scheme 3) also affected the mechanical response. Our study presented a more comprehensive evaluation of tension-compression switching strategies in the GOH model by not only presenting mathematical clarification of continuum theory and also linking with their implementations in commonly used simulation tools as used by the modeler.

This study filled an important knowledge gap in brain computational biomechanics by systematically examining the influence of tension-compression switches in the GOH model for WM modeling. Despite the wide usage of the GOH model for brain material modelling (see the review by Wright and Daphalapurkar [76] and introduction of current study), only one early study [27], to the best of the author’s knowledge, noted that different switching criteria were used in the GOH model to simulate the WM as a fiber-reinforced anisotropic structure. That study also speculated that such discrepancies could affect the mechanical response of brain tissue with dependency on the level of anisotropy, although detailed verification was not presented. The current study conducted a comprehensive comparison to highlight the strain difference among three candidate switching schemes, including strain-time history curves of representative elements (Fig 4), qualitative visualization of WM elements with fibers in compression (Fig 5) and quantitative differences of strain peaks at the element level (MPS) and the fiber level (MFS) (Fig 6) in a concussive impact, and statistical analysis of peaking strain responses at a group level (Fig 7). The observed inter-scheme differences could be explained by equations 2-4, wherein both the switching parameter and the treatment of fibers in compression differed among schemes. Our multi-aspect protocol for cross-scheme strain comparison, along with the theoretical elaboration in Section 2.3 and the extensive body of continuum mechanics literature mentioned earlier, collectively highlighted the importance of tension-compression switches in the GOH model for WM anisotropy. It was also worth noting that all three switching schemes were implemented via user-defined material subroutines, representing a technical advancement that enabled focused comparisons within a unified computational framework.

This study found that *Ē*_*α*_>0 does not necessarily require the fiber to be in tension (*Ī*_4*α*_>1) (see the *Ī*_4*α*_-*Ē*_*α*_ curve of scheme 1 at the compression regime in Fig 2). Such a finding highlighted a fundamental limitation of adopting *Ē*_*α*_ as the switching parameter, as it failed to distinguish between tensile and compressive fiber states. From the aspect of material modelling, it might be possible that the *Ē*_*α*_ switch was not activated even when fibers were kinematically compressed, violating the underlying modeling assumption that axonal fibers contributed only in tension and not in compression. Such behavior introduced an artificial contribution of fibers in compression, which could lead to spurious stiffening and distorting the predicted anisotropic response. From the aspect of injury predictability, fiber strain (or alternatively termed as axonal strain, tract-oriented strain, tract-oriented normal strain in the literature [28, 77-79]) was often used as an brain injury metric [76]. An inappropriate switch might lead to either overestimation or underestimation of fiber recruitment, thereby biasing injury predictions. Although this issue has been repeatedly articulated in prior theoretical and computational studies [27, 48, 74, 75, 80], its practical implications appeared to have been insufficiently recognized. For example, Abaqus software continues to employ *Ē*_*α*_ as the switching variable in its built-in GOH formulation and has been widely used for brain anisotropic modelling [26, 35, 36, 38, 42]. Similar issues were also noted several other WM material modelling studies based on user-defined material [28, 37, 39-41]. It is advised that caution should be exercised when interpreting the anisotropic responses from these studies. Together with the previous work [27, 48, 74], the current study advocates the adoption of *Ī*_4*α*_ as the switching parameter, as was originally proposed by Gasser, et al. [34].

In addition to the paradoxical switching schemes, the tension-compression switch itself constitutes a fundamental assumption of the GOH model. However, to the best of the author’s knowledge, direct experimental evidence of compression-induced buckling at the level of WM fiber/axon/neuron appeared lacking. For example, Reiter, et al. [81] subjected histologically stained fresh human and porcine brain tissue specimens to compressive loadings at the tissue scale and simultaneously observed the resulting displacement of embedded microstructures through an inverse microscope. It was clearly noted therein that the blood vessel buckled during the loading, but when the compressive loading was along the main axon direction, no clear axonal buckling was reported. In parallel, several *in vitro* studies reported the compressive axial strain endured by neurons [82-84], implying no evident buckling was observed. The absence of direct experimental confirmation of the tension-compression switch made it infeasible to recommend a specific implementation (e.g., scheme 2 vs. scheme 3 or even other alternatives). This knowledge gap might also partially explain the large body of literature that assumed the fiber/axon/neuron endured loadings in compression [85-87]. It should be noted that, as critically analyzed by Horgan and Murphy [88], the tension-compression switch hypothesis was also challenged in the modelling of other biological tissues, such as muscle [89], tendon [90], and arterial tissue [91, 92].

At an indirect level, several recent brain tissue experiments did provide partial support for the tension-compression switch hypothesis. For example, Reiter, et al. [19] applied multimodal, multidirectional loading to the tissue sample dissected from the human corpus callosum. When imposing compressive loading to the sample, significantly higher stresses were noted when loaded transverse to the fibers compared to loading in the fiber direction. In tension, an opposite trend was noted with significantly higher maximum stresses along the fiber direction than transverse to it. Such results could imply that the nerve fibers in the corpus callosum contributed to tissue strength in tension but not in compression. Nevertheless, in the same experiment by Reiter, et al. [19], the human medulla oblongata tissue samples showed no clear relationship between loading direction and fiber orientation observed, indicating the tension-compression switch might be region-dependent. Taken all together, there is a clear need for more comprehensive test protocols to fully characterize the three-dimensional mechanical responses and relate the experimental data with underlying microstructural architectures from medical imaging and histological quantification, directly verifying or rejecting the tension-compression hypothesis.

Similar to the strategies in other FE human head models [26, 27, 30, 38, 42-44, 86, 93], the current study employed the GOH model to simulate the brain tissue in the KTH model as an anisotropic structure, which was supported by a substantial body of literature reporting that the brain tissue exhibited mechanical anisotropy [19, 94, 95]. Nevertheless, the authors acknowledged that a considerable number of experimental studies instead reported an isotropic mechanical behavior of the brain tissue. For example, Jin, et al. [96] found the mechanical stiffness of WM in tension and compression showed no significant dependency on the fiber direction. Budday, et al. [97] reported that the isotropic one-term Ogden model was capable of simultaneously representing the hyperelastic behavior under combined shear, compression, and tension loadings. Indeed, the majority of existing FE human brain models simulated the brain as an isotropic structure [98-103]. The authors acknowledge that both material characterization and constitutive modelling of brain tissue are active research areas, and the mechanical anisotropy of WM remains to be a matter of debate. Relevant discussions on this topic are elaborated in several reviews [104, 105] and our previous publications [73, 106] and are not reiterated herein.

Several additional limitations, other than the few highlighted above, should be acknowledged. First, the current study employed an FE model with relatively coarse meshes to improve computational efficacy and the GOH model was implemented at the element level with a single embedded fiber orientation, representing significant losses of orientation information. Nevertheless, the observed mechanical effects of the tension-compression switch were supported by the theoretical explanation (equations 2-4) and were therefore not expected to depend strongly on the specific FE discretization. Second, experimental studies on the guinea pig optic nerves [107] and the developing chick embryo spinal cord [108] reported a gradual coupling between the undulated axons and the surrounding glial cells secondary to mechanical stretching, but similar data on human axonal fibers was lacking. Considering the inter-species variation, extrapolation of the results derived from animal experiments to human subjects needs further verification [109]. The current study thereby assumed the fiber was straight in its undeformed configuration. Finally, the current study considered the fiber dispersion using the general structure tensor approach proposed by Gasser, et al. [34]. Alternative methods, such as the angular integration approach [110-112], have been proposed in recent years but have not yet been applied to brain tissue modelling, representing an another interesting direction for future work.

## 5. Conclusions

The current study systematically evaluated the role of tension-compression switch formulations in the GOH model for anisotropic modelling of WM. The results demonstrated that both the choice of switching parameter (*Ē*_*α*_ vs. *Ī*_4*α*_) and the treatment of fiber contributions in compression (0 vs. *κ*(*Ī*_1_ − 3)) led to significant differences in predicted brain strain responses, highlighting the sensitivity of model predictions to these implementation choices. The Green-Lagrange strain-like quantity *Ē*_*α*_ was shown to be an unreliable indicator of fiber stretch, advocating the adoption of *Ī*_4*α*_ as the switching parameter. This study highlighted the need for direct verification of the tension-compression switch hypothesis in axonal fibers, as well as improved experimental characterization towards a converged understanding of the mechanical anisotropy of brain tissue.

## Acknowledgments

This research has received funding from KTH Royal Institute of Technology (Stockholm, Sweden), Swedish Research Council (VR-2024-05848, and VR-2024-02782), and FFI (Strategic Vehicle Research and Innovation), project numbers 2023-00753 and 2024-03635, funded by Vinnova, the Swedish Transport Administration, the Swedish Energy Agency, and the industrial partners. The content of this article is solely the responsibility of the authors and does not necessarily represent the official views of neither funding agencies. We’d also like to acknowledge Professors Svein Kleven and T. Christian Gasser at KTH, Dr. Richard Davies at EDRMedeso for their stimulating discussion on the brain constitutive modelling and practical advice of software implementation. The computational simulations were enabled by resources provided by the National Academic Infrastructure for Supercomputing in Sweden (NAISS) at the center for High Performance Computing (PDC) partially funded by the Swedish Research Council through grant agreement no. 2022-06725. The authors declare that they have no known competing financial interests or personal relationships that could have appeared to influence the work reported in this paper.

## Author contributions

**Chengbin Li:** Methodology, Software, Investigation, Visualization, Data curation Writing – original draft, review & editing; **Zhou Zhou:** Conceptualization, Methodology, Software, Writing – review & editing; Funding acquisition.

